# *Aggregatibacter actinomycetemcomitans* LtxA hijacks endocytic trafficking pathways in human lymphocytes

**DOI:** 10.1101/769778

**Authors:** Edward T Lally, Kathleen Boesze-Battaglia, Anuradha Dhingra, Nestor M Gomez, Claire H Mitchell, Alexander Giannakakis, Syed A Fahim, Roland Benz, Nataliya Balashova

## Abstract

Leukotoxin (LtxA) from oral pathogen *Aggregatibacter actinomycetemcomitans* is a secreted membrane-damaging protein. LtxA is internalized by β2 integrin LFA-1 (CD11a/CD18) expressing leukocytes and ultimately causes cell death; however toxin localization in the host cell is poorly understood and these studies fill this void. We investigated LtxA trafficking using multi-fluor confocal imaging, flow cytometry and Rab5 knockdown in human T lymphocyte Jurkat cells. Planar lipid bilayers were used to characterize LtxA pore-forming activity at different pH. Our results demonstrate that LtxA/LFA-1 complex gains an access to the cytosol of Jurkat cells without evidence of plasma membrane damage utilizing dynamin-dependent and clathrin-independent mechanism. Upon internalization LtxA follows the LFA-1 endocytic trafficking pathways as identified by co-localization experiments with endosomal and lysosomal markers (Rab5, Rab11A, Rab7, and Lamp2) and CD11a. Knockdown of Rab5a resulted in loss of susceptibility of Jurkat cells to LtxA cytotoxicity suggesting that late events of LtxA endocytic trafficking are required for toxicity. The toxin trafficking via the degradation endocytic pathway may culminate in delivery of the protein to lysosomes or its accumulation in Rab11A-dependent recycling endosomes. The ability of LtxA to form pores at acidic pH may result in permeabilization of the endosomal and lysosomal membranes.

## 1. INTRODUCTION

The RTX (<u>R</u>epeats in <u>T</u>o<u>X</u>in) toxins are membrane-damaging proteins secreted by several Gram-negative bacteria [1]. The organisms producing these proteins are important human and animal pathogens implicating the toxins role in the bacteria virulence. RTX-toxins common features are the unique mode of export across the bacterial envelope via the type I secretion system employing an uncleaved C-terminal recognition signal [2–4] and the characteristic nonapeptide glycine-and aspartate-rich repeats binding Ca^2+^ ions [5,6]. The toxins are modified with fatty acid moieties attached to internal lysine residues which is an unusual characteristic for bacterial proteins [7–10]. RTX toxins can be divided into three groups: (i) broadly cytolytic RTX hemolysins, (ii) species-specific RTX leukotoxins, and (iii) large multifunctional autoprocessing RTX toxins (MARTX)[1]. RTX leukotoxins exhibit a narrow cell type and species specificity due to cell-specific binding through protein receptors of the β_2_ integrin family[1]. The β_2_ integrins are expressed on the surface of leukocytes and share a common β_2_ subunit, CD18, which is combined with either one of the unique α chains, α_L_ (CD11a), α_M_ (CD11b), α_X_ (CD11c), or α_D_ (CD11d)[11].

*Aggregatibacter actinomycetemcomitans* (*Aa*), a facultative anaerobe and common inhabitant of the human aerodigestive tract, causes localized aggressive periodontitis (LAP) [12]. Localized aggressive periodontitis (LAP) is a rapidly progressing periodontal disease that results in loss of tooth attachment and alveolar bone destruction in adolescents. If left untreated in teenagers, the infection will lead to the loss of the permanent first molars and central incisors [13]. *Aa* produces several virulence determinants that contribute to either bacterial colonization or destruction of the periodontium. The pivotal virulence factor of *Aa* is an RTX leukotoxin, LtxA, that kills both human innate and adaptive immune cells [14]. *Aa* isolated from LAP patients predominantly belongs to a single clone JP2 [15], which is characterized by increased LtxA production, implicating a role for LtxA in disease development [16]. Analysis of primary LtxA sequence consisting of 1055 amino acids predicts four LtxA domains [17]. The hydrophobic domain encompassing residues 1-420 is followed by the central domain (residues 421-730) that contains two internal lysine residues (K^562^ and K^687^) that are the sites of post-translational acylation, required for LtxA activation [9]. The repeat domain (residues 731-900) contains the characteristic repeated amino acid sequence of the RTX family with the C-terminal domain (residues 901-1055) hypothesized to play a role in secretion [17].

LtxA toxicity requires the presence of the β_2_ integrin LFA-1(LFA-1, CD11a/CD18 or α_L_/β_2_) [18] and cholesterol enriched membrane domains [19,20]. LFA-1 is a native ligand for intercellular adhesion molecule (ICAM-1) located on vascular endothelial cells [21]. In immunocytes LFA-1/ICAM-1 binding is one of the molecular mechanisms for leukocyte adhesion and migration to the site of infection [22]. LFA-1 is constantly internalized and then rapidly recycled back to the plasma membrane through vesicular transport [23,24] using the “long-loop” of recycling involving GTPase Rab11A [25]. Additionally, LFA-1 activity toward the components of the extracellular matrix is regulated by the ability of these receptors to switch between active and inactive conformations [21]. Cholesterol is essential for LtxA association with plasma membrane of human lymphocytes [19] and human monocytes [20] [26]. Binding to cholesterol is mediated by a cholesterol recognition amino acid consensus (CRAC) motif, in the N-terminal region of the toxin [19].

Recent findings suggest that recirculating and resident memory T cells in gingival tissue play an important role in maintenance of periodontal homeostasis [27]. In an experimental rat periodontal disease model, antigen-specific CD4 T lymphocytes were required for bone resorption [28]. In our study Jurkat cell line, subclone Jn.9, served as model to investigate LtxA uptake and trafficking in T lymphocytes. These cells express cell-surface LFA-1 and are susceptible to LtxA induced toxicity [29]. Initial interaction of LtxA with the host membrane elevates cytosolic Ca^2+^ independent of the toxin and LFA-1. Ca2+ elevation involves activation of calpain, talin cleavage, and subsequent clustering of LFA-1 in lipid rafts on the membrane [26]. In the proposed mechanism of the LtxA/LFA-1 interaction, LtxA binds to the extracellular domains of LFA-1 subunits, CD11a and CD18. The toxin then transverses the cell membrane, binds to the cytoplasmic tails of LFA-1, and causes activation of LFA-1 [30]. Following from the results of the liposomal study LtxA adopts a U-shaped conformation in the membrane, with the N- and C-terminal domains residing outside of the membrane [31].

After binding to the LFA-1 subunits, LtxA has been quickly internalized into the cytosol where it was found in vesicular structures [30]. The pathway of intracellular LtxA trafficking has not been investigated. Endocytosed toxins initially accumulate in endosomes, where they may take advantage of the acidic environment within the vesicles to form, or contribute to membrane damage in order to translocate into the cytosol. Since LtxA binds to LFA-1, the possibility is that LtxA could be using an integrin trafficking pathway to gain access to the target cell cytosol. Here we examined the components of the cytosol of LtxA-treated cells for co-localization of the toxin and the CD11a subunit of LFA-1 with different organelle markers. LtxA association with endosomal and lysosomal markers suggests a receptor mediated endocytic process that may culminate in delivery of the toxin to lysosomes. Additionally, the toxin can be redirected to the plasma membrane due to the LFA-1 receptor Rab11a-mediated recycling. This study provides new insight into convergent mechanism of LFA-1 and LtxA trafficking, and the ability of LtxA to function in acidic environments.

## 2. RESULTS

### 2.1. LtxA does not damage host cell membrane when enters the cell

The membrane damaging properties of LtxA have been documented [32,33]. Therefore, the first question we asked was whether initial steps of LtxA interaction with the cells result in the plasma membrane damage. The green-fluorescent impermeable nucleic acid stain YO-PRO®-1 (630 kDA) is used to detect early membrane damage and permeates cells immediately after membrane destabilization [34]. We performed flow cytometry analysis of LtxA-treated cells to determine YO-PRO®-1 internalization to indicate plasma membrane damage. The YO-PRO®-1 membrane permeabilization assay showed no evidence of plasma membrane damage in LtxA treated Jn.9 cells within first 4 h of the treatment (Fig. 1). However, our flow cytometry data demonstrate that 20 nM LtxA-DY488 become internalized with Jn.9 cells as early as 15 min after the toxin was added and its internalization steadily increased over time. Our results indicate that LtxA is quickly internalized by Jn.9 cells but the toxin does not rupture the plasma membrane when it enters the host cells.

**Figure 1.**
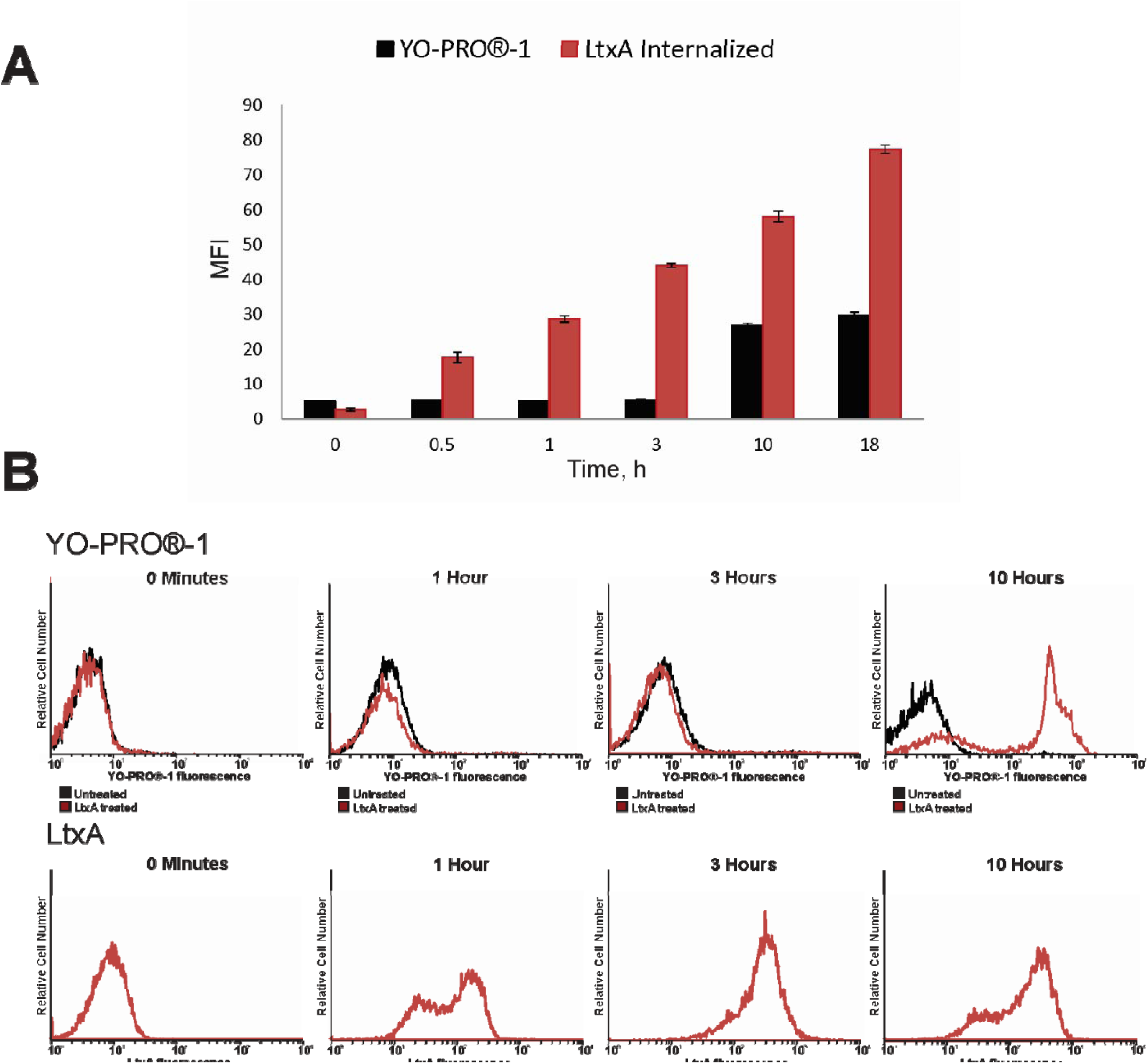
Flow cytometry analysis of YO-PRO®-1 and LtxA internalization with Jn.9 cells. Flow cytometry analysis was used to detect YO-PRO®-1 and LtxA inside of Jn.9 cells over time. Cells (1 × 10^6^) were incubated with 10 µM YO-PRO®-1 alone or in presence of 20 nM LtxA. Another set of cells was treated with 20 nM LtxA-DY488 at different times. The extracellular fluorescence of the cells was quenched with 0.025% trypan blue [30] and the intracellular fluorescence was determined. **A.** Uptake of YO-PRO®-1 (black) and internalization of LtxA-DY488 (red) at different times presented as mean fluorescence intensity (MFI) of the entire cell population. The data shown is the result of three independent experiments. Error bars indicate ±SEM. **B.** Top: flow cytometry histograms showing YO-PRO®-1 dye uptake by LtxA treated cells (red line) v.s. the dye uptake by untreated Jn.9 cells (black line) at different times. Bottom: flow cytometry histograms showing LtxA-DY488 internalized with Jn.9 cells. The data shown is representative of three independent experiments.

### 2.2. LtxA uptake is diminished by dynamin inhibitors

We used a set of chemical and pharmacological inhibitors of endocytosis (Table 1) to define the mechanism of the toxin uptake by Jn.9 cells. Fluorescent-labeled toxin internalization was significantly reduced in cells pre-treated with dynamin-inhibitors. To confirm the efficiency or specificity of selected dynamin inhibitors concentrations, the internalization of transferrin labeled with Alexa Fluor®647 was followed using confocal microscopy. Cells pretreated with 10 µM Dynasore and 5 µM Dynole 34-2, which block GTPase activity of dynamin [35] [36], for 20 min were less susceptible to LtxA. However, the inhibitors affecting clathrin-mediated endocytic pathway such as potassium depleted medium and 5 µM Pitstop 2 did not change activity of LtxA on Jn.9 cells. This suggests that LtxA internalization in Jn.9 cells is dynamin-dependent (Fig. 2).

**Table 1.** Chemical inhibition of LtxA uptake.

| Compound | Mode of action | Effect |
| --- | --- | --- |
| 10 µM Dynasore* | Blocks GTPase activity of dynamin [35] | Inhibits internalization/activity |
| 10 µM Dynole 34-2* | Blocks GTPase activity of dynamin [36] | Inhibits internalization/activity |
| 10 µM Dynole 31-2* | Inactive derivative of Dynole 34-2 | No effect |
| 10 µM Pitstop 2* | Interferes with binding of proteins to<br>the N-terminal domain of clathrin [81] | No effect |
| Metyl-β-cyclodextrin | Removes cholesterol from plasma membrane [82] | Inhibits activity [26] |
| K<sup>+</sup>-depletion | Inhibits clathrin mediated endocytosis [83] | Inhibits internalization |

\*To measure LtxA activity inhibition, Jn.9 cells were preincubated with 10 µM inhibitors for 20 min in serum free medium. The chemicals stocks were prepared in DMSO and were added in the 1 µl volume to 1 ml of cells. No adverse effect of DMSO alone on Jn.9 cells was observed.

**Figure 2.**
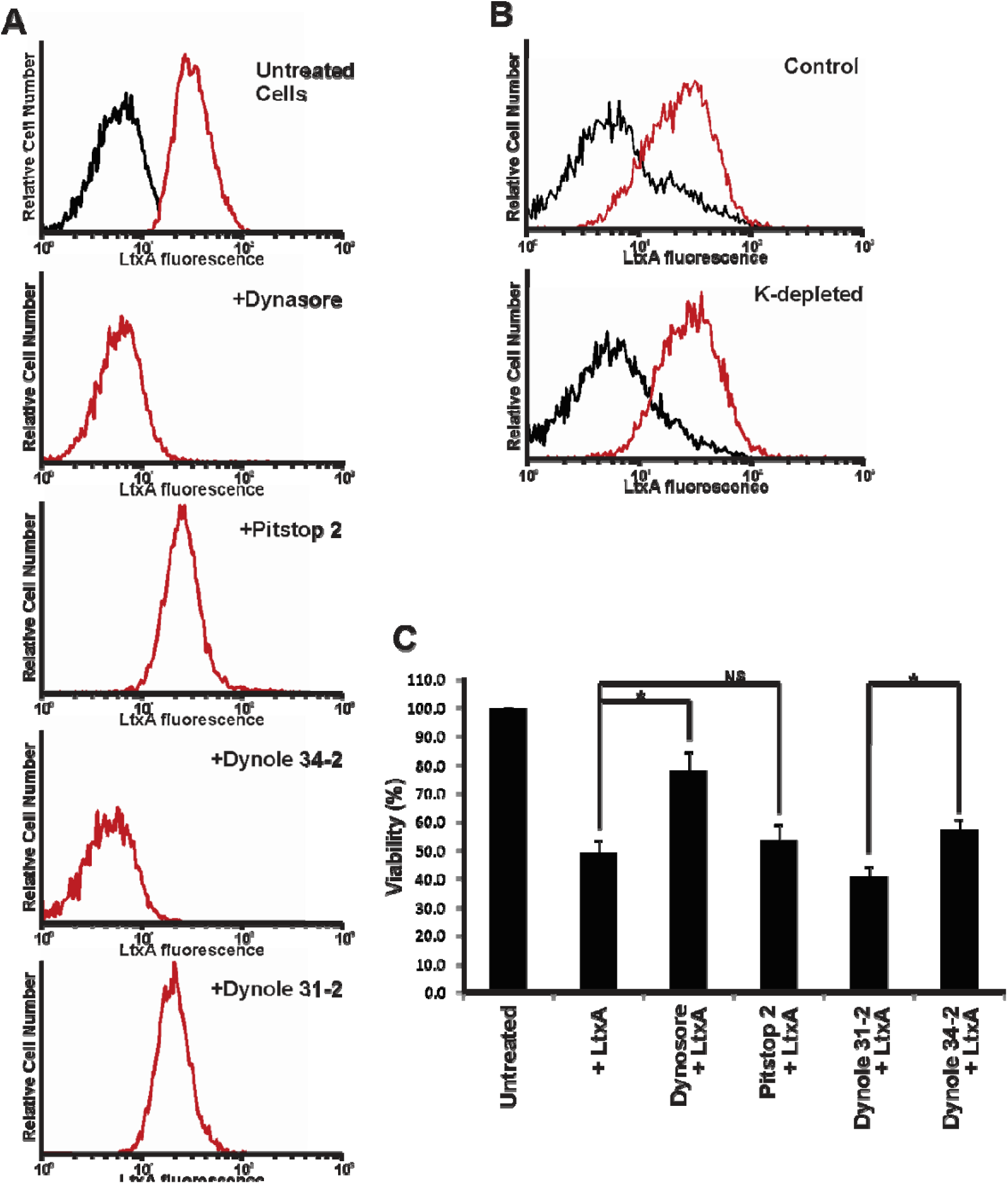
Effect of endocytosis inhibitors on LtxA internalization and activity. **A. Flow cytometry analysis of LtxA-DY488 internalization with Jn.9 cells pretreated with chemical inhibitors**. Jn.9 cells (1 × 10^6^) were preincubated with endocytosis inhibitors for 20 min, and then were treated with 20 nM LtxA-DY488 for 30 min at 37 °C. The extracellular fluorescence of the cells was quenched (0.025% trypan blue) [78,79] and intracellular cell fluorescence (red peak) was determined by flow cytometry analysis. No residual fluorescence was detected in 0.1% Triton X-100 permeabilized cells after the trypan blue treatment. Untreated cells (black) served as a negative control. Representative flow cytometry histograms are shown. **B. Flow cytometry analysis of LtxA internalization with Jn.9 cells in potassium-depleted medium**. Jn.9 Cells (1 × 10^6^ cells) were incubated in K^+^-containing (top) or K^+^-depleted buffer (bottom), and then 20 nM LtxA-DY488 was added for 30 min. Flow cytometry analysis to determine the amount of internalized toxin (red peak) was performed as described in Fig. 2A. Untreated cells (black) served as a negative control. Representative flow cytometry histograms are shown. **C. LtxA toxicity on Jn.9 cells**. To measure LtxA activity inhibition, Jn.9 cells were preincubated with 10 µM inhibitors for 20 min in serum free medium at 37 °C, and then 10 nM LtxA was added and the cells were incubated for 18 h. The cell viability was measured by the trypan blue assay. The cells treated with specific chemical served as a negative control. LtxA killing efficiency was adjusted accordingly. No adverse effect of DMSO alone on Jn.9 cells was observed. Error bars indicate ±SEM, *p ≤ 0.05 compared with corresponding chemical treated cells. The data shown is the result of three independent experiments.

### 2.3. LtxA and CD11a are found in early and recycling endosomes

Jn.9 cells treated with fluorescent-labeled LtxA for 30 min were used to performing immunocytochemistry experiments with endocytic pathway markers including GTPase Rab5 and Rab11a. The co-distribution of LtxA and LFA-1 heterodimer components on the surface of target cell membranes suggests that LtxA could gain access to the cytosol as individual LtxA molecules or as part of an LtxA/LFA-1 complex. In our imaging studies LtxA was found in vesicular structures after entry into Jn.9 cells. Our immunocytochemistry studies demonstrated co-localization of LtxA, early endosome membrane protein Rab5a and CD11a suggesting toxin uptake through receptor-mediated endocytosis. Fig. 3A and Fig. 1S-A show confocal images of Jn.9 cells with co-localization of LtxA, CD11a, and early endosomal marker Rab5a after treatment of the cells with LtxA-DY650 for 30 min at 37 °C. We found co-localization of CD11a and LtxA with recycling endosomal marker Rab11a in recycling endosomes. LFA-1 are exocytosed via GTPase Rab11A-mediated recycling [37] a process that involves trafficking through the perinuclear recycling compartment (PNRC), before reaching the plasma membrane. Co-localization of CD11a and LtxA with recycling endosomal marker Rab11a in recycling endosomes (Fig. 3B and Fig. 1S-B) suggests that after entering the early endosome a significant amount of LtxA is redirected back to the membrane in LFA-1 recycling turnover. Alternatively, release of LtxA into PNRC can provide access to the nuclear membrane for LtxA. Indeed in our imaging studies we often observed the toxin surrounding nuclei (Fig. 2S).

**FIGURE 3.**
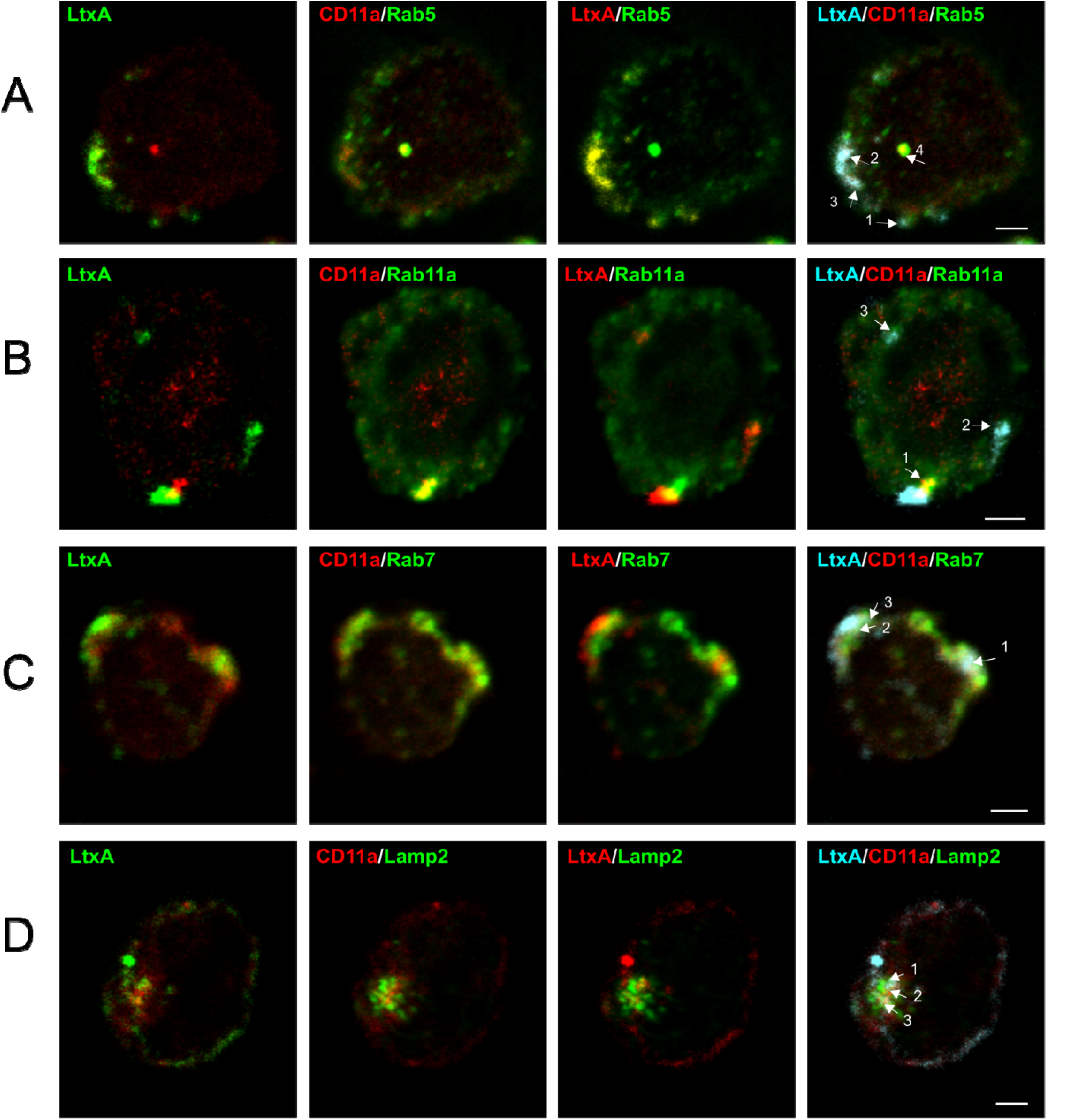
Confocal imaging of LtxA trafficking in Jn.9 cells. Confocal images showing localization of LtxA, CD11a, endosome or lysosome markers in Jn. 9 cells after treatment with the toxin for 30-120 min at 37 °C. LtxA-DY650 is pseudo colored in green, CD11a detected with mouse anti-CD11a antibody conjugated with Alexa Fluor™594 is shown in red, and endosomal/lysosomal markers detected through goat-anti rabbit antibody conjugated with DyLight™ 488 are shown in green.. Arrows demonstrate the areas of co-localization. No significant background fluorescence in unstained Jn.9 cells was detected. Last panel shows triple staining with LtxA (cyan), Cd11a (red), and Rab5/Rab11/Rab7 or Lamp2 (green). Representative images are shown. Scale bars = 2μm. **A.** Localization of LtxA, CD11a and Rab5 after treatment of the cells with the toxin for 30 min. Pearson’s coefficients for LtxA/Rab5 co-distribution are: 1=0.60, 2=0.85 and 3=0.79; for LtxA/CD11a co-distribution: 1=0.65, 2=1.2 and 3=0.8; for CD11a/Rab5 co-distribution: 1=0.65, 2=0.82, 3=0.72 and 4=0.96. **B.** Localization of LtxA, CD11a and Rab11a after treatment of the cells with the toxin for 30 min. Pearson’s coefficients for LtxA/Rab11a co-distribution are: 1=0.81, for LtxA/CD11a co-distribution: 1=0.8, 2=0.95 and 3=0.8; for CD11a/Rab11a co-distribution: 1=0.93. **C.** Localization of LtxA, CD11a and Rab7 after treatment of the cells with the toxin for 1 h. Pearson’s coefficients for LtxA/Rab7 co-distribution are: 1=0.87, 2=0.74 and 3=0.72; for LtxA/CD11a co-distribution: 1=0.92, 2=0.59 and 3=0.77; for CD11a/Rab7 co-distribution: 1=0.52, 2=0.53, and 3=0.94. **D.** Localization of LtxA, CD11a and Lamp2 after treatment of the cells with the toxin for 2 h. Pearson’s coefficients for LtxA/Lamp2 co-distribution are: 1=0.78, 2=0.65 and 3=0.72; for LtxA/CD11a co-distribution: 1=0.06, 2=-0.34 and 3=-0.02; for CD11a/Lamp2 co-distribution: 1=0.32, 2=0.25, 3=0.11.

### 2.4. Rab5 siRNA knockdown limits LtxA toxicity

Irrespective of routes of internalization endocytic cargoes are trafficked to early endosomes, where Rab5 GTPases is the key player in subsequent trafficking events [38]. We investigated the impact of Rab5 downregulation on LtxA uptake and toxicity on cells (Fig. 4). Western blot analysis 24 h after transfection confirmed that Rab5 was significantly downregulated (≥90%) in Jn.9 cells compare to scrambled siRNA transfected cells. When transfected cells were treated with 2nM LtxA for 18 h the toxic effect of the toxin on Rab5 downregulated cells was 30% less than on control cells (Fig. 4A). Internalization of LtxA was analyzed by flow cytometry after 30 min of treatment with 20 nM LtxA-DY488. No significant variations in the amount of internal fluorescence were detected in cells transfected with Rab5 siRNA (MFI 7.4) and cells using scrambled siRNA (MFI 6.9) (Fig. 4B). Our results suggest that abolishment of Rab5 function does not affect LtxA internalization or cytotoxicity.

**FIGURE 4.**
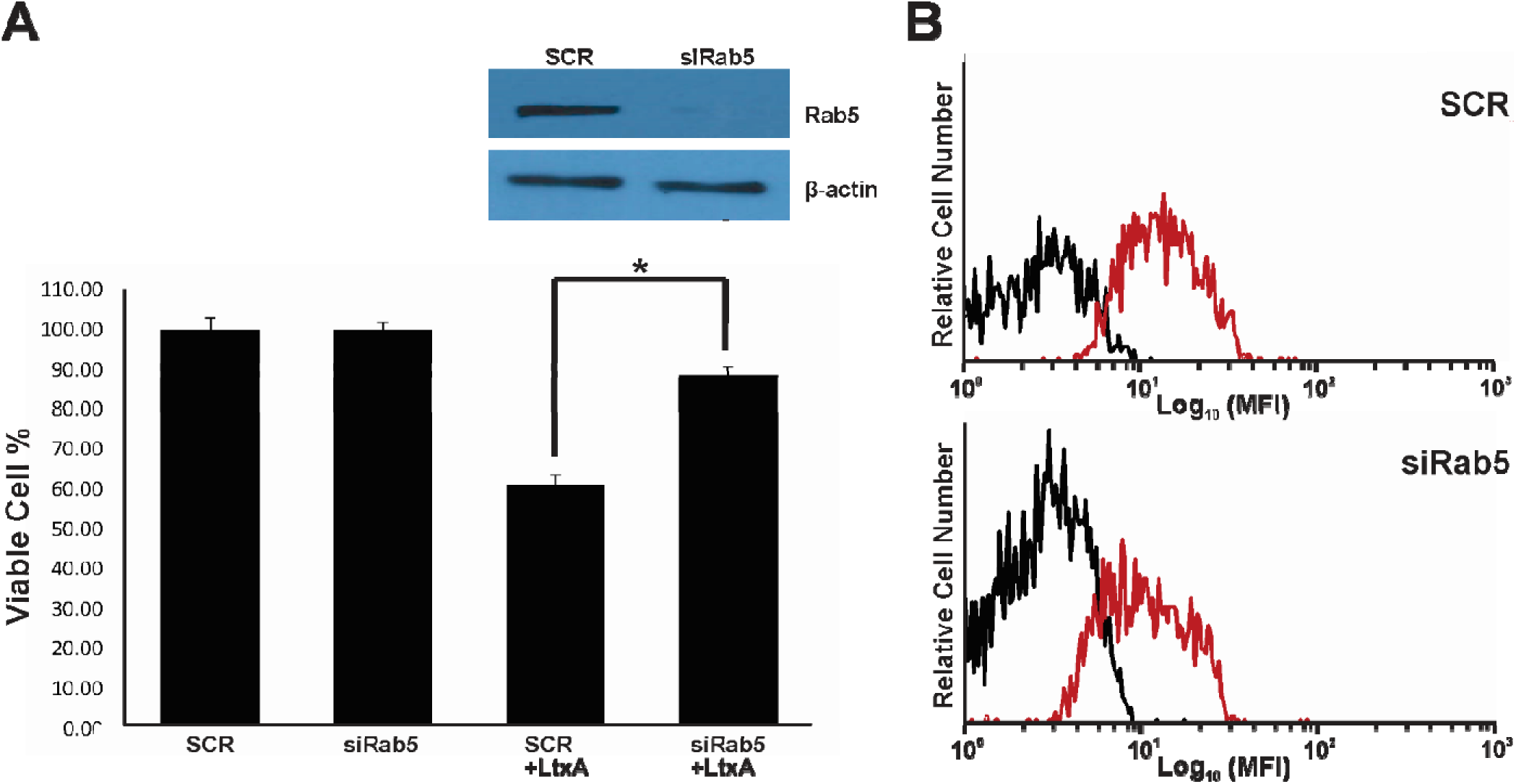
Modulation of Rab5 function in Jn.9 cells. **A.** Jn.9 cells (1 × 10^6^ cells) were transfected with siRNA control (SCR) or with siRNA against Rab5 and collected 24 h post-transfection for Rab5a expression analysis by Western blotting. The cell viability testing was performed by trepan blue assay after 18 h of treatment with 20 nM LtxA. A representative expression of Rab5a protein was shown for 24 h of siRNA treatment. The Rab5a protein expression (inset) was normalized to β-actin expression. Error bars indicate ±SEM, *p ≤ 0.05 compared with siRNA SCR-treated cells.The experiment was performed three independent times. **B.** Jn.9 cells (1 × 10^6^ cells) were transfected with siRNA control (SCR) or with siRNA against Rab5a were collected 24 h post-transfection and then treated with 20 nM LtxA-DY488 for 30 min at 37 °C. The extracellular fluorescence of the cells was quenched (0.025% trypan blue) [78,79] and intracellular cell fluorescence (red peak) was determined by flow cytometry analysis. No residual fluorescence was detected in 0.1% Triton X-100 permeabilized cells after the trypan blue treatment. Untreated (black) served as a negative control. Representative flow cytometry histograms are shown.

### 2.5. LtxA and CD11a are found in late endosomes and lysosomes

At later time points Jn.9 cells treated with fluorescent-labeled LtxA were used in immunocytochemistry experiments with the endocytic pathway markers GTPases Rab7 and Lamp2. After 1 h of treatment with LtxA-DY650, LtxA associated with the late endosome membrane protein Rab7 and CD11a (Fig. 3C and Fig. 1S-C). Co-localization of LtxA with lysosomal marker Lamp2 after 2 h of treatment with LtxA-DY650 indicated that the toxin trafficking culminates in its delivery to the lysosomes, where LtxA was found separated from CD11a (Fig. 3D and Fig. 1S-D).

### 2.6. LtxA causes lysosomal damage in Jn.9 cells

We detected LtxA in Jn.9 lysosomes and therefore we wanted to see whether LtxA was able to cause lysosomal damage in the cells. We have probed the effect of LtxA on lysosomal integrity in the cells using lysosomal dye, LysoTracker® Red DND-99, and followed changes in lysosomal properties of the cells using live imaging after addition of 20 nM LtxA to the cells by live cell confocal microscopy. The loss of the fluorescence intensity in LtxA treated cells, but not in untreated cells, indicated that LtxA caused damage of lysosome. No changes in lysosomal fluorescence were detected within the first 90 min of treatment and about 15% decrease of LysoTracker® Red DND-99 staining intensity was identified in Jn.9 cells after 2 h of treatment (Fig. 5), which may indicate lysosomal damage due to lysosomal membrane permeabilization or lysosome alkalization.

**FIGURE 5.**
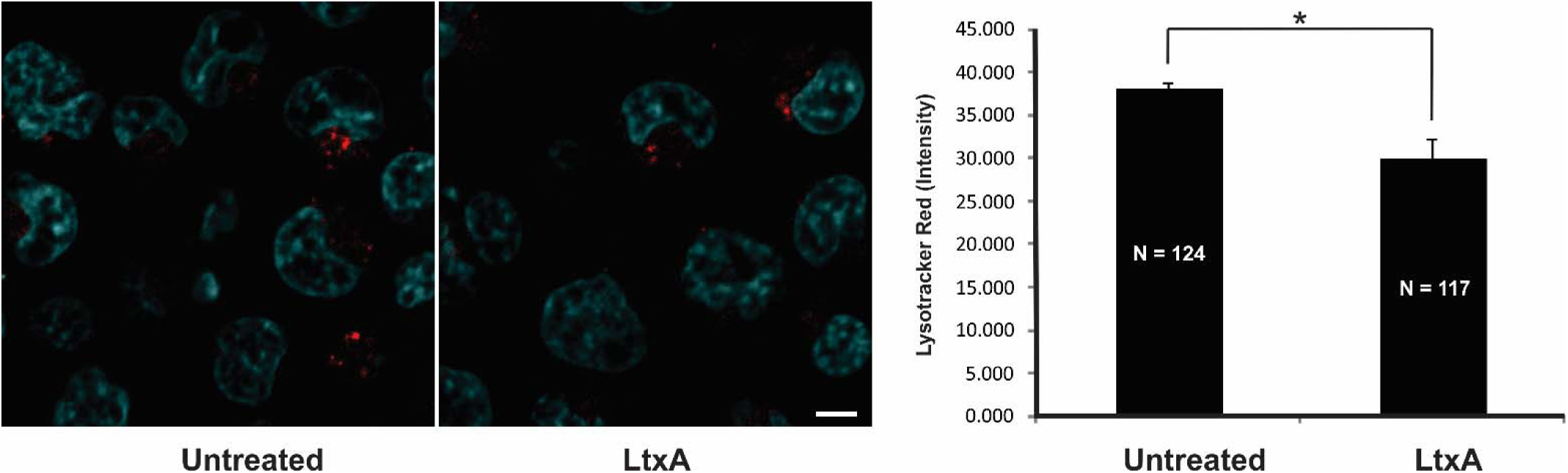
Decrease in lysosomal pH after treatment with LtxA. Lysosomes in Jn.9 cells were stained with LysoTracker^®^ Red DND-99 (red) and nuclei were stained with Hoechst 33342 (pseudo colored in cyan). The confocal images of cells collected after treatment with 20 nM LtxA (2 h) or untreated are shown on the left. Average red fluorescence intensity per cell calculated in N (number) cells is shown on the right. Error bars indicate ±SEM. *P ≤ 0.05 compared with untreated cells lysosomal intensity. Representative images are shown and are the results of three independent experiments. Scale bar = 10 μm.

### 2.7. LtxA is active in lipid bilayer membranes at low pH

Channel formation by LtxA was studied in detail at neutral pH [33,39]. In lipid bilayer membranes formed of asolection LtxA forms cation selective channels with a single-channel conductance of approximately 1.2 nS in 1 M KCl (pH 6.0) [39]. Since LtxA is found in endocytic vesicles, we asked whether LtxA is also able to form ion-permeable channels at acidic pH. To address this, we performed lipid bilayer experiments with wildtype LtxA at different pH-values ranging from pH 3.5 to pH 10.0. LtxA formed ion-permeable channels in 1 M KCl solutions under all these conditions (pH 3.5, 4.7, 7.5, 8.5 and 10.0). However, because the membranes became very fragile at very low and very high pH (3.7 and 10.0) it was not possible to record too many single-channel events under these conditions. At the other pH-values the membranes were rather stable and a sufficient number of single-channel events could be recorded in the experiments. Fig. 6 shows a single channel recording of LtxA in 1 M KCl, 10 mM MES-KOH, pH 4.7. The channel had a somewhat reduced lifetime at this pH as compared with that a neutral pH [39]. Fig. 3B and C shows a histogram obtained from 47 LtxA channels recorded under these conditions. A fit of the histogram with a Gaussian function yielded an average single-channel conductance of 1.1 ± 0.3 nS somewhat smaller than that at pH 6.0 (G = 1.2 ± 0.3 nS) [39]. Again we found that the single-channel distribution was quite broad, similar to the conditions at pH 6.0 (Fig. 6, Table 2).

**Table 2.** Influence of the aqueous pH on the conductance of channels formed by LtxA.

| Salt and Buffer | pH | $G^* \pm SD$ (nS) | Number of events n |
| --- | --- | --- | --- |
| 1 M KCl, 10 mM MES-KOH | 3.7 | $1.0 \pm 0.21$ | 12 |
| 1 M KCl, 10 mM MES-KOH | 4.7 | $1.1 \pm 0.31$ | 47 |
| 1 M KCl, 10 mM MES-KOH | 6.0 | $1.2 \pm 0.30$ | 95 |
| 1 M KCl, 10 mM Tris-HCl | 7.5 | $1.2 \pm 0.24$ | 39 |
| 1 M KCl, 10 mM Tris-HCl | 8.5 | $1.3 \pm 0.29$ | 53 |
| 1 M KCl, 10 mM Tris-HCl | 10 | $1.2 \pm 0.26$ | 14 |

\*The LtxA conductance ( $G \pm \text{variance}/SD$ ) in each 1M KCl solution was either taken from Gaussian distributions (see Fig. 3) or directly from the statistics of single-channel data (n number of single events). To analyze the conductance in each case n channels were reconstituted in asolectin/n-decane membranes at 20 mV voltage, $t=20^\circ\text{C}$ . The number of events analyzed at pH 3.7 and 10 was low due to instability of the lipid bilayers at extreme pH.

**Figure 6.**
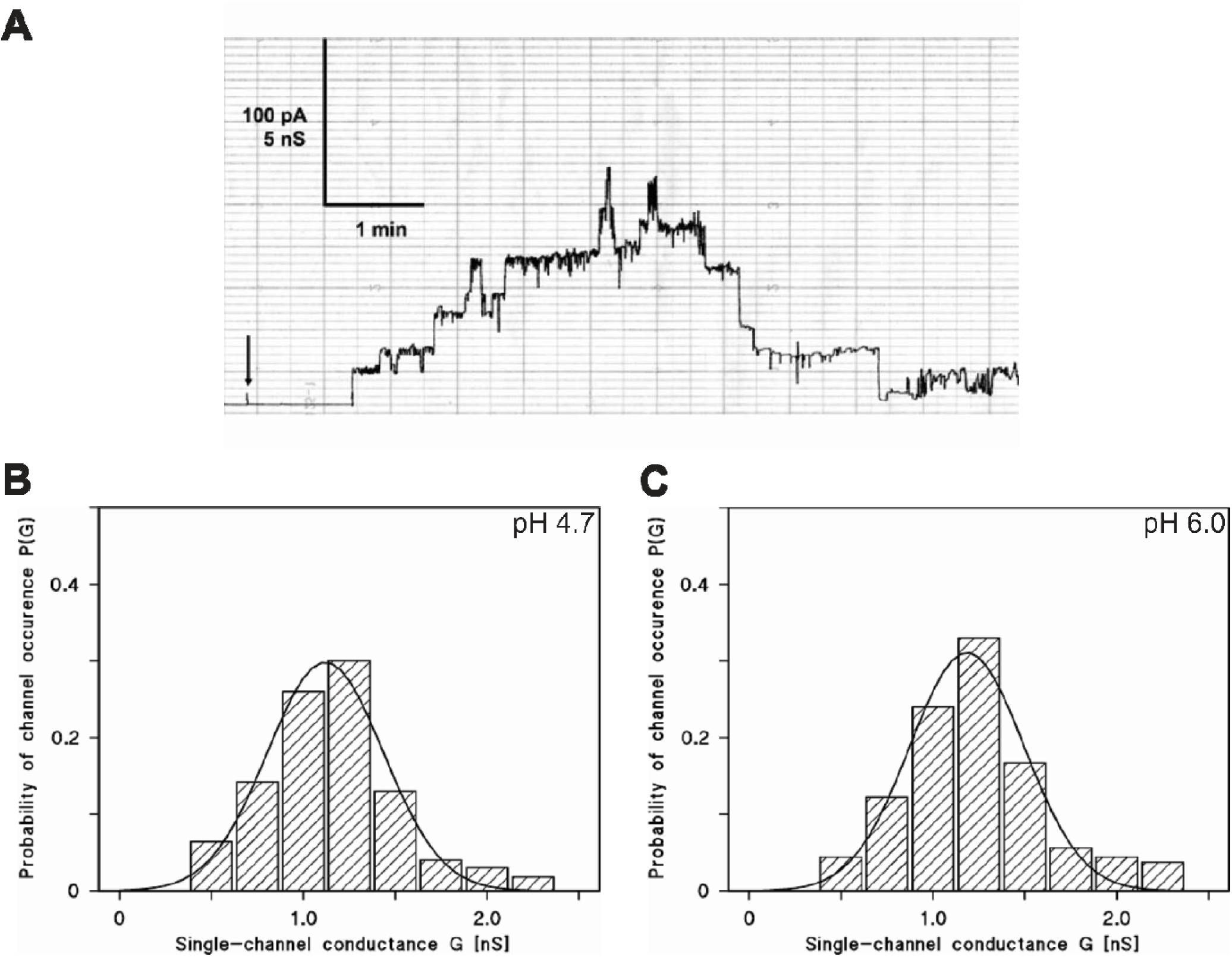
Pore forming activity of LtxA in asolectin/*n*-decane membranes at different pH. **A.** Single-channel recording of LtxA in an asolectin/n-decane membrane at pH 4.7. Current recording of an asolectin/n-decane membrane performed in the presence of 10 nM LtxA added to the cis-side of the membrane. The aqueous phase contained 1 M KCl, 10 mM MES-KOH, pH 4.7. The applied membrane potential was 20 mV at the cis-side (indicated by an arrow), t=20°C. **B.** Histogram of the probability P(G) for the occurrence of a given conductivity unit observed for LtxA with membranes formed of 1% asolectin dissolved in *n*-decane in a salt solution at pH 4.7. The histogram was calculated by dividing the number of fluctuations with a given conductance unit by the total number of conductance fluctuations. The average conductance of all conductance steps was 1.1 ± 0.31 nS for 47 conductance steps derived from 9 individual membranes. The value was calculated from a Gaussian distribution of all conductance fluctuations (solid line). The aqueous phase contained 1 M KCl, 10 mM MES-KOH, pH 4.7 and 10 nM LtxA, the applied membrane potential was 20 mV, t=20°C. **C.** Histogram of the probability P(G) for the occurrence of a given conductivity unit observed for LtxA with membranes formed of 1% asolectin dissolved in n-decane in a salt solution at pH 6.0. The average conductance of all conductance steps was 1.20 ± 0.31 nS for 95 conductance steps derived from 17 individual membranes. The aqueous phase contained 1 M KCl, 10 mM MES, pH 6.0 and about 10 nM LtxA, the applied membrane potential was 20 mV, t=20°C.

We studied also the effect of high pH on channel formation mediated by LtxA. Ion-permeable channels were also observed at these conditions. The average single channel conductance at pH 7.5, 8.5 and 10 is shown in Table 2. The influence of the aqueous pH was rather small on the conductance of the LtxA channel despite a possible shift of the selectivity of the LtxA channel from slightly cation selective at pH 6.0 to a higher selectivity for potassium ions over chloride.

## 3. DISCUSSION

Leukocytes need to quickly transmigrate from blood vessels into tissues upon inflammation or infection. An essential mechanism regulating this process is the subcellular trafficking of adhesion molecules, primarily integrins [40]. Integrins undergo constant endo/exocytic turnover necessary for the dynamic regulation of cell adhesion. Bacterial toxins have developed a number of schemes to cross the membrane in order to enter the cell. LtxA evolved the strategy to target specifically β_2_ integrin LFA-1 on leukocytes’surface [18]. This binding is required for the toxin internalization [30].

We here report that LtxA is delivered to the lysosomes of Jn.9 cells through endocytic trafficking. Historically, endocytic pathways are classified as either clathrin-dependent or clathrin-independent. The large GTPase dynamin [41] is thought to be directly involved in pinching off endocytic vesicles from the plasma membrane. The key players in the formation of clathrin coated vesicles are dynamin [41] and adaptor proteins [42]. The studies with *Mannheimia haemolytica* LktA, another RTX leukotoxin, show that LktA is internalized in a dynamin-2 and clathrin-dependent manner [43]. The following LktA-trafficking events involve the toxin binding to the mitochondria and interaction with cyclophilin D, a mitochondrial chaperone protein, in bovine lymphoblastoid cells [44].

Our results indicate that LtxA enters Jn.9 cells using clathrin-independent mechanism (or predominantly uses this pathway). Our results correlate with the finding that LFA-1 is internalized through a clathrin-independent cholesterol-dependent pathway and this process is essential for cell migration [45]. In this scenario non-clathrin-coated lipid raft microdomains form 50–100 nm flask-shaped vesicular invaginations of the plasma membrane regions rich in lipid rafts [46]. Lipid-raft dependent endocytosis was shown to be dynamin-dependent [47] and may involve caveolae formation, which require participation of caveolin-1. Okadaic acid, a potent inhibitor of specific protein phosphatases, causing the removal of caveola from the cell surface [48]. Thus, we hypothesize that LtxA/LFA-1 is endocytosed through caveolae-mediated endocytosis. In agreement with that, we identified that Jn.9 cells express caveolin-1 on their surface and treatment of the cells with okadaic acid inhibited LtxA toxicity (data not shown).

Bacterial toxins often hijack existing endocytic trafficking pathways [49,50] to deliver active protein to subcellular targets. The small GTPases Rab are key regulators of intracellular membrane trafficking and exist in an inactive GDP-bound form and an active GTP-bound form [51]. The co-localization experiments with Rab5, Rab7, Lamp2 revealed that LtxA can follow the degradation pathway process that culminate in delivery of the toxin to lysosomes. Rab5 localizes to early endosomes where it is involved in the recruitment of Rab7 and the maturation of these compartments to late endosomes [52]. Impaired Rab5 function affects endo- and exocytosis rates and in opposite, Rab5 overexpression increases the release efficacy [53]. Therefore, termination of Rab5 function blocks movement of proteins downstream endocytic pathway. Downregulation of Rab5 decreased LtxA toxicity suggesting that further toxin trafficking is required for intoxication by LtxA. LFA-1 undergoes endocytic recycling through long-Rab11 dependent pathway with transitional step at PNRC. While some LtxA follows LFA-1 in its recycling turnover, a portion of LtxA is separated from LFA-1 and the toxin proceeds to late endosomes and lysosomes. Proposed model of LtxA trafficking in lymphocytes is shown on Fig. 7.

**FIGURE 7.**
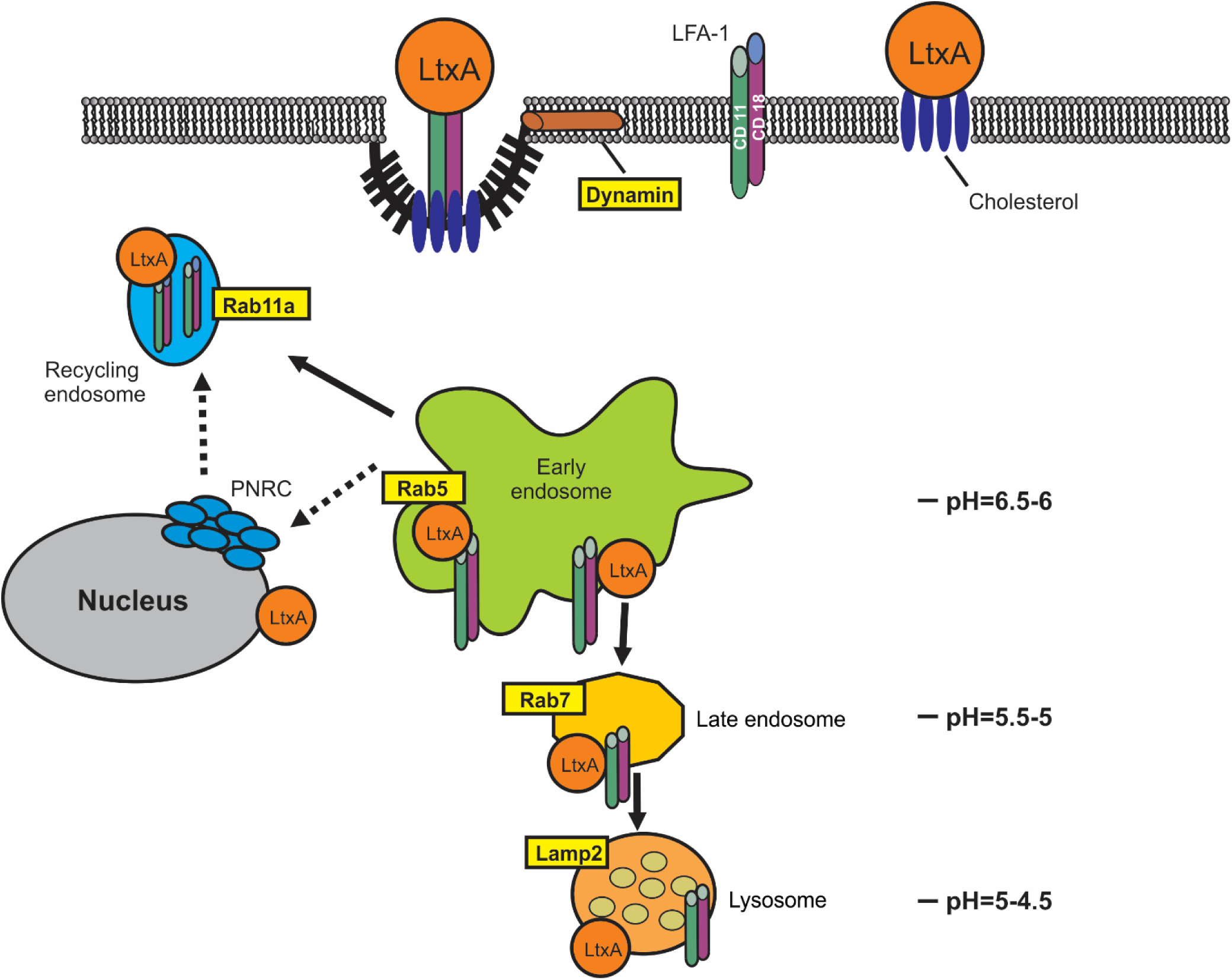
Proposed mechanism of LtxA entry and trafficking in human lymphocytes. LtxA binds to cholesterol and LFA-1 on the surface of Jn.9 cell. LtxA/LFA-1 complexes internalization is dynamin dependent. Internalized LtxA/LFA-1 complexes are quickly transported to early endosomes. The small GTPase Rab5 regulates membrane docking and fusion in the early endocytic pathway. Interruption of Rab5 expression in Jn.9 cells results abolishment of the LtxA activity. LFA-1 undergoes endocytic recycling through long-Rab11 dependent pathway with transitional step at PNRC [25]. While some LtxA follows LFA-1 in its recycling turnover, a portion of LtxA is separated from LFA-1 and the toxin proceeds to late endosomes and lysosomes. The ability of LtxA to damage lipid membranes at low pH may cause endocytic vesicles and lysosomal rupture and release of the toxin to the cytosol.

Interaction between integrins and their β-integrin ligands typically lead to enhanced cell survival and several immunological changes [54,55]. Our experiment using cell impermeable dye, YO-PRO®-1, serves to demonstrate that LtxA gains access to the Jn.9 cell cytosol without evidence of plasma membrane damage. Our study and others suggest that LtxA could accumulate in lysosomes and alter lysosomal pH [56,57]. Damage of lysosomes by LtxA in human and rat monocytes cells [58,59] and in human erythroleukemia cells [58] were reported. In our previous study, treatment with 100 ng/ml LtxA leaded to cytosol acidification in K562 cells expressing LFA-1, presumably due to leakage of lysosomal content, as was identified using a pH sensitive indicator pHrodo^®^. This process correlated with the disappearance of lysosomes in the cytosol as determined by both acridine orange and LysoTracker® Red DND-99 staining. Similarly, using LysoTracker® Red DND-99 dye, lysosomal damage was detected in malignant monocytes (THP-1 cells) as early as 15 min after treatment with LtxA and reached 70% after 2 h of treatment (unpublished data). In these cells LtxA was shown to localize to the lysosome where it induces active cathepsin D release [59]. Here we demonstrate that LtxA causes changes in lysosomal pH in T lymphocytes, however to a leser extent. The pore-forming properties of LtxA are well established [33,60]. Therefore, we propose that LtxA can cause permeabilization of lysosomal membrane, and possibly other intracellular organelles after the toxin is released from lysosomes. Alternatively, LtxA can accumulate in lysosomes altering pHL. The toxin molecule possesses a hydrophobic regions and is modified with acyl residues. Fat molecules may accumulate in cell lysosomes increasing pHL and disrupt normal lysosomal function as in lysosomal storage disorder (LSDs)[61].

LtxA was reported to cause different cellular responses leading to cell death in LFA-1 expressing cells. Kelk et al. reported that LtxA lyses healthy monocytes by activation of inflammatory caspase 1 and causes release of IL-1b and IL-18. In contrast to myeloid cells, LtxA uses “slow mode” of lymphocyte killing. Killing of malignant lymphocytes requires Fas receptors and caspase 8 in both T and B lymphocytes [62]. In B lymphocytes (JY cells), LtxA caused loss of mitochondrial membrane potential, cytochrome c release, reactive oxygen species release, and activation of caspases 3,7,9 [26]. One possible explanation for the cell death mechanism induced by LtxA is the degree of lysosomal damage caused by the toxin in cell. The extent of lysosomal rupture will determine morphological outcomes following lysosomal membrane permeabilization. Extensive lysosomal damage may lead to inevitable necrotic cell death, while less extensive detriment of lysosomes may induce several apoptotic pathways, which can be attenuated by inhibition of lysosomal proteases (cathepsins) [63–65].

The planar lipid bilayer assay is a highly sensitive method that allows characterization of membrane damaging activity of RTX-toxins in different physical conditions [66]. A current model proposes that RTX-toxins form cation-selective channels with diameter 0.6 – 2.6 nm in artificial membranes formed of lipid mixtures such as asolectin/n-decane membrane [39]. It was demonstrated that membrane-damaging activity of LtxA in artificial bilayers did not require the presence of the receptor [67]. In endocytic pathway, subsequent acidification may initiate proteolysis and conformational changes resulting in the ability of toxins and viruses to cross the endocytic vesicle membrane since drugs that interfere with the endosomal pH are able to block the infection [68,69]. In this study we used this method to observe and compare pore formation of LtxA at different pH. We demonstrated that LtxA is functional in acidic pH found in endocytic vesicles and lysosomes, which may result in their damage. RTX toxins are intrinsically disordered proteins, therefore changes in pH may affect their secondary structure and consequently change their activity [70]. Further investigation is required to our understanding of the intracellular events leading to LtxA-induced cytolysis.

## 4. MATERIALS AND METHODS

### 4.1. Antibodies and chemicals

The following primary antibodies were used; CD11a Alexa Fluor™ 594 clone HI111 (Biolegend, San Diego, CA), rabbit polyclonal anti-Rab5, anti-Rab11A, anti-Rab7, or anti-Lamp2 antibody (Abcam, Cambridge, UK), anti-beta-actin antibody (AnaSpec, Fremont, CA) (1:1000), and anti-LtxA monoclonal antibody 107A3A3 [71] in hybridoma supernatants (1:10 dilution). The following secondary antibodies were used: goat anti-rabbit IgG Alexa Fluor^®^ 488 (1:1000); horseradish peroxidase (HRP)-conjugated goat anti-mouse IgG (Fc) or (HRP)-goat anti-rabbit (Pierce, Rockford, IL) (1:10,000). Transferrin labeled with Alexa Fluor®647 was from Invitrogen (Waltham, MA, USA). Dynamin inhibitor Dynole 34-2 and its inactive control, Dynole 31-2, were purchased from SigmaAldrich (St. Louis, MO), Dynasore and Pitstop 2 (Abcam, Cambridge, UK). The inhibitors were used in the following concentrations: 10 μM Dynole 34-2; 10 μM Dynole 31-2; 10 μM Dynasore; 5 μM Pitstop 2.

### 4.2. Cell culture

Jn.9, a subclone of Jurkat cells [72] was utilized in this study. The cells were cultivated in RPMI 1640 medium containing 10% FBS, 0.1 mM MEM non-essential amino acids, 1x MEM vitamin solution, and 2 mM L-glutamine, and either 0.5 μg/mL gentamicin at 37 ºC under 5% CO_2_.

### 4.3. LtxA purification and labeling

*Aa* strain JP2 [73] was grown on solid AAGM medium [74] for 48 h at 37 °C in 10% CO_2_ atmosphere. The colony was then inoculated in 1.5 L of liquid AAGM medium and the culture was incubated for 18 h. The toxin was purified from cell culture supernatants as described previously [75]. Purified LtxA was labeled with DyLight™ 650 (LtxA-DY650) or DyLight™ 488 (LtxA-DY488) using DyLight™ Amine-Reactive dyes (Pierce) according to previously published protocol [76].

### 4.4. Immunofluorescence

For LtxA trafficking studies, 1×10^6^ of Jn.9 cells were incubated with 20 nM LtxA-DY650 for 15 min to 2 h at 37 ºC in the growth medium. The cells were then washed with PBS, fixed with 2% paraformaldehyde for 10 min, washed twice with PBS, and permeabilized with 0.2% Triton X-100 for 20 min. The cells were subsequently blocked with 5% BSA for 30 min at 37 °C, incubated with primary antibody for 18 h at 4 ºC, washed, and incubated with secondary antibody conjugated to Alexa Fluor 488 for 1 h at 37 °C. The nuclei were stained with 1 μg/ml Hoechst 33342 (Molecular Probes™, Eugene, OR) for 15 min at 37 °C. Samples were mounted in Cytoseal mounting medium (Electron Microscopy Sciences, Hatfield, PA) and images captured with a Nikon A1R laser scanning confocal microscope with a PLAN APO VC 60 × water (NA 1.2) objective at 18°C. Data were analyzed using Nikon Elements AR 4.30.01 software; for co-distribution analyzes, the Pearson’s’ coefficient was at least 0.55, and analysis included maximum intensity projection and standard LUT adjustment.

For lysosomal staining, Jn.9 cells were treated with 20 nM LtxA for 2 h at 37°C in the growth medium. Then the cells were washed with The Live Cell Imaging Solution (LCIS) (Molecular Probes™, Eugene, OR) and treated with 100 nM LysoTracker^®^ Red DND-99 (Life Technologies, Carlsbad, CA) and 1 μg/ml Hoechst 33342 for 15 min at 37 °C. After another wash with LCIS the cells were placed to attach for 20 min in ibiTreat 60 μ-dishes (Ibidi, Madison, WI) coated with poly-L-lysine (Sigma St. Louis, MO), then they were washed again and covered with LCIS. The cells were examined using a Nikon A1R laser scanning confocal microscope with a 60× water objective (NA 1.2). Approximately 100 cells per image were analyzed in each experiment to identify the mean fluorescent intensity per cell by sorting non-saturated areas in three combined Z planes collected for each image.

### 4.5. Inhibitors

Chemicals stocks were prepared in DMSO and were added in the 1 µl volume to 1 ml of cells. To measure LtxA activity inhibition, Jn.9 cells (1 × 10^6^ cells) were pre-incubated with 5-10 µM inhibitors for 20 min in the serum free medium at 37 °C. For cytotoxicity evaluation, the cells were treated with 2 nM LtxA and the cell viability was evaluated as described in the “Cytotoxicity assay” section. The toxin internalization assay was performed as described in the “Flow cytometry” section. The cells treated with specific inhibitors served as a negative control. The effect of K^+^-depleted medium was evaluated using previously published protocol [77].

### 4.6. Flow cytometry

YO-PRO®-1 internalization was investigated using Membrane permeability/dead cell apoptosis kit (Invitrogen, Carlsbad, CA) according to the manufacture’s protocol. To detect internalized LtxA, Jn.9 cells (1 × 10^6^ cells/run) were incubated with 20 nM LtxA-DY488 for 30 min at 37 °C in PBS supplemented with 2% FBS, washed with PBS, and total cell-associated fluorescence was analyzed. To quench the extracellular fluorescence, LtxA-DY488 treated cells were incubated with 0.025% trypan blue (Sigma, St. Louis, MO) for 20 min as described previously [78,79]. To quench the intracellular fluorescence cells were permeabilized using 0.1% Triton X-100 (SigmaArdrich, St. Louis, MO) for 10 min prior to 0.025% trypan blue treatment. Fluorescence was measured using a BD LSR II flow cytometer (BD Biosciences). Ten thousand events were recorded per sample, and the mean fluorescence intensity (MFI) values were determined using WinList™7.0 software (Verity Software House). To quantitate the intracellular fluorescence, MFI values of cells pretreated with trypan blue were subtracted from the MFI values of total cell-associated LtxA-AF™488 fluorescence. No residual fluorescence was detected in 0.1% Triton X-100 permeabilized cells after the trypan blue treatment. Samples that were not treated with LtxA-DY488 served as a control.

### 4.7. Protein analyses

The protein concentration was determined by absorption at 280 nm on A1 NanoDrop spectrophotometer (Thermo Fisher Scientific, Waltham, MA). Proteins were resolved on 4 to 20% SDS-PAGE and visualized by staining with GelCode blue stain reagent (Pierce, Rockford, IL). The Western blot analysis was performed as described previously [66].

### 4.8. siRNA

The validated Silencer® Select siRNA targeting human Rab5a (ID s11678) and Silencer^®^ Select Negative Control #2 siRNA (catalog# 4390846) were synthesized by Life Technology (Carlsbad, CA). Jn.9 cells were transfected with lipofectamine 2000 (Life Technologies, Carlsbad, CA) according to the manufacturer’s instructions. For each transfection, 5 µl of the 20 µM siRNA stocks were added to 400 µl of Jn.9 cells grown to 90% confluency. Rab5a levels were confirmed by Western blot analysis 24 h after transfection.

### 4.9. Cytotoxicity assay

For toxicity tests 2-20 nM LtxA was added to 1 × 10^6^ Jn.9 cells in growth medium and incubated for 18 h at 37 °C. The cell membrane permeability was determined with trypan blue assay using Vi-cell Cell Viability Analyzer (Beckman Coulter, Miami, FL). All reactions were run in duplicate; the assay was performed three independent times. Untreated cells were used as controls.

### 4.10 Planar lipid bilayers

Lipid bilayer measurements are described previously in detail [80]. In brief, A Teflon chamber divided into two 5 mL compartments that are connected by a small circular hole with a surface area of about 0.4 mm^2^ were filled with 1 M KCl, 10 mM MES, pH 6.0. Black lipid bilayer membranes were obtained by painting onto the hole solutions of 1% (w/v) asolectin (phospholipids from soybean, Sigma-Aldrich) or diphytanoyl phosphatidylcholine (DiphPC; Avanti Polar Lipids, Alabaster, AL) in *n*-decane. All salts were analytical grade and the temperature was maintained at 20°C during all experiments. The current across the membrane was measured with a pair of Ag/AgCl electrodes with salt bridges switched in series with a voltage source and a highly sensitive current amplifier Keithley 427 (Keithley Instruments, INC. Cleveland, OH). The amplified signal was recorded by a strip chart recorder (Rikadenki Electronics GmbH, Freiburg, Germany).

### 4.11 Statistical Analysis

The statistical analyses were performed using either Student’s *t* test or one-way analysis of variance using SigmaPlot^®^ (Systat Software, Inc. Chicago, IL). The following statistical criteria were applied: *p* < 0.001, *p* < 0.05, and *p* < 0.01.

## Supplementary Materials

Figure S1: “Confocal imaging of LtxA trafficking in Jn.9 cells”; Figure S2: “Localization of LtxA around nuclear membrane of Jn.9 cells”.

## Author Contributions

Conceptualization, Edward T Lally; Data curation, Nestor M Gomez and Nataliya Balashova; Formal analysis, Nestor M Gomez, Anuradha Dhingra, Roland Benz and Nataliya Balashova; Funding acquisition, Edward T Lally; Investigation, Nestor M Gomez, Alexander Giannakakis, Syed A Fahim and Roland Benz; Methodology, Kathleen Boesze-Battaglia, Claire H Mitchell and Roland Benz; Project administration, Nataliya Balashova; Software, Anuradha Dhingra; Supervision, Nataliya Balashova; Writing – review & editing, Kathleen Boesze-Battaglia and Claire H Mitchell.

## Funding

This work was supported by the United States National Institute of Health grants R01DE009517 (ETL and NB), R01DE022465 (KBB).

## Acknowledgements

The authors thank Juan Reyes-Reveles (JRRDesign Inc.) for technical assistance.

---

**FIGURE 1S.**
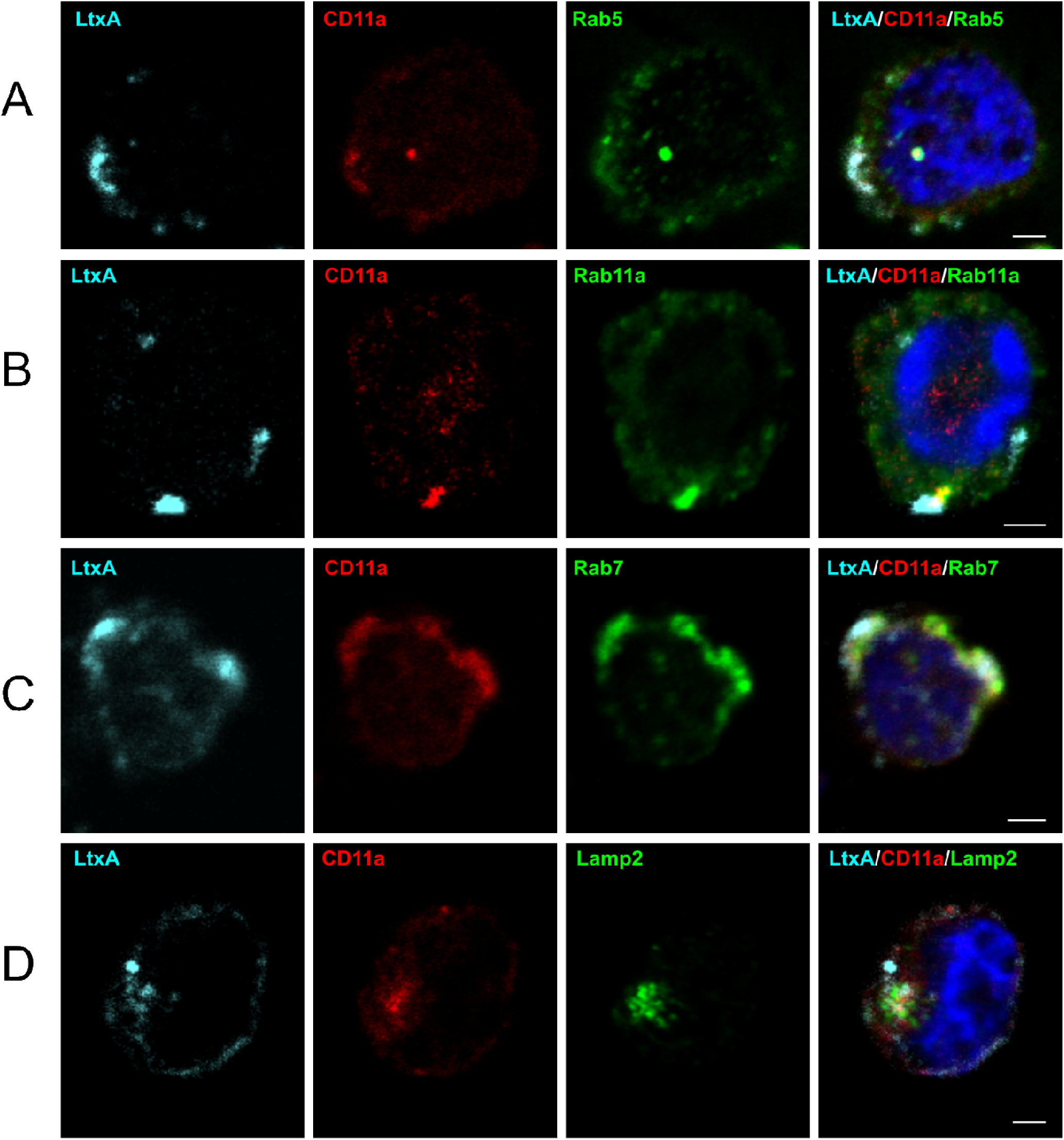
Confocal imaging of LtxA trafficking in Jn.9 cells. Confocal images showing localization of LtxA, CD11a, endosome or lysosome markers in Jn. 9 cells after treatment with the toxin for 30-120 min at 37 °C. LtxA-DY650 is shown in cyan, CD11a detected with mouse anti-CD11a antibody conjugated with Alexa Fluor™594 is shown in red, and endosomal/lysosomal markers detected through goat-anti rabbit antibody conjugated with DyLight™ 488 are shown in green. Cell nuclei were stained with Hoechst 33342 (blue). No significant background fluorescence in unstained Jn.9 cells was detected. Representative images are shown. Scale bars = 2μm. **A**. Localization of LtxA, CD11a and Rab5 after treatment of the cells with the toxin for 30 min. **B**. Localization of LtxA, CD11a and Rab11A after treatment of the cells with the toxin for 30 min. **C**. Localization of LtxA, CD11a and Rab7 after treatment of the cells with the toxin for 1 h. **D**. Localization of LtxA, CD11a and Lamp2 after treatment of the cells with the toxin for 2 h.

**FIGURE 2S.**
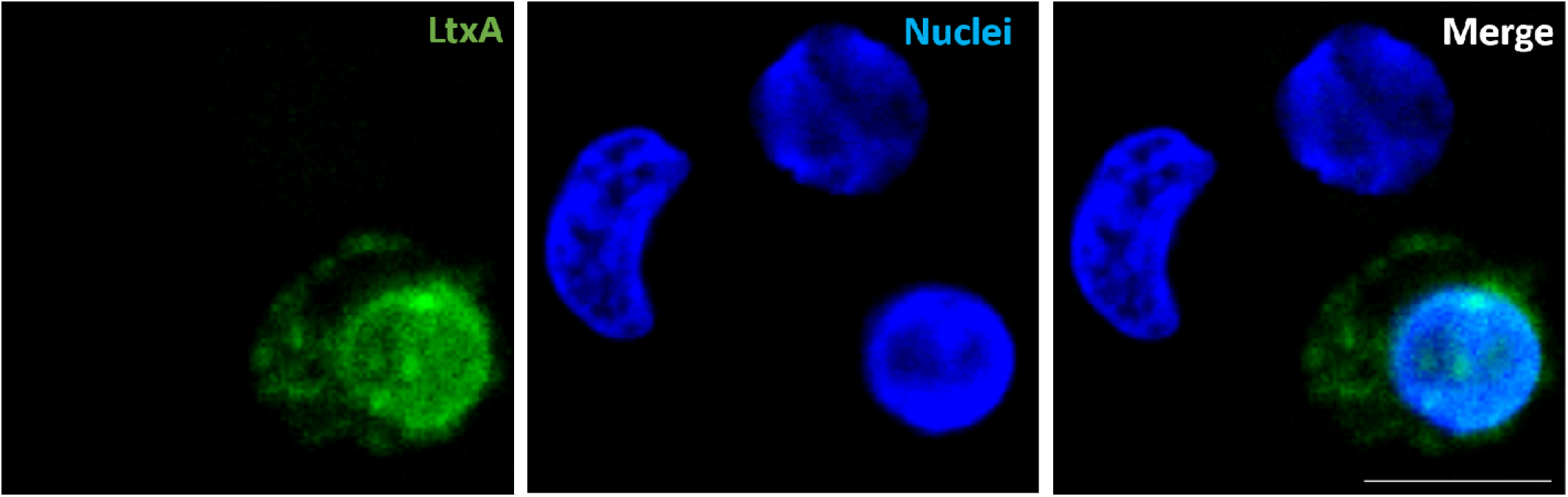
Localization of LtxA around nuclear membrane of Jn.9 cells. Confocal images of Jn.9 cells showing localization of LtxA-DY650 after treatment of the cells with the toxin for 2 h at 37 °C. The immunostaining was performed as described in Materials and Methods. LtxA-DY650 (pseudo colored in green) and cell nuclei were stained with Hoechst 33342 (blue). No significant green background fluorescence in unstained Jn.9 cells was detected. Representative images are shown. Scale bar = 10μm.

## LITERATURE CITED

1. Linhartova, I.; Bumba, L.; Masin, J.; Basler, M.; Osicka, R.; Kamanova, J.; Prochazkova, K.; Adkins, I.; Hejnova-Holubova, J.; Sadilkova, L., et al. RTX proteins: a highly diverse family secreted by a common mechanism. FEMS Microbiol. Rev. 2010, 34, 1076–1112, doi:10.1111/j.1574-6976.2010.00231.x.

2. Goni, F.M.; Ostolaza, H. E. coli alpha-hemolysin: a membrane-active protein toxin. Braz. J. Med. Biol. Res. 1998, 31, 1019–1034.

3. Ostolaza, H.; Soloaga, A.; Goni, F.M. The binding of divalent cations to Escherichia coli alpha-haemolysin. Eur. J. Biochem. 1995, 228, 39–44.

4. Soloaga, A.; Veiga, M.P.; Garcia-Segura, L.M.; Ostolaza, H.; Brasseur, R.; Goni, F.M. Insertion of Escherichia coli alpha-haemolysin in lipid bilayers as a non-transmembrane integral protein: prediction and experiment. Mol. Microbiol. 1999, 31, 1013–1024.

5. Boehm, D.F.; Welch, R.A.; Snyder, I.S. Domains of Escherichia coli hemolysin (HlyA) involved in binding of calcium and erythrocyte membranes. Infect. Immun. 1990, 58, 1959–1964.

6. Coote, J.G. Structural and functional relationships among the RTX toxin determinants of gram-negative bacteria. FEMS Microbiol. Rev. 1992, 8, 137–161.

7. Stanley, P.; Packman, L.C.; Koronakis, V.; Hughes, C. Fatty acylation of two internal lysine residues required for the toxic activity of Escherichia coli hemolysin. Science 1994, 266, 1992–1996.

8. Stanley, P.; Koronakis, V.; Hughes, C. Acylation of Escherichia coli hemolysin: a unique protein lipidation mechanism underlying toxin function. Microbiol. Mol. Biol. Rev. 1998, 62, 309–333.

9. Balashova, N.V.; Shah, C.; Patel, J.K.; Megalla, S.; Kachlany, S.C. Aggregatibacter actinomycetemcomitans LtxC is required for leukotoxin activity and initial interaction between toxin and host cells. Gene 2009, 443, 42–47, doi:10.1016/j.gene.2009.05.002.

10. Osickova, A.; Balashova, N.; Masin, J.; Sulc, M.; Roderova, J.; Wald, T.; Brown, A.C.; Koufos, E.; Chang, E.H.; Giannakakis, A., et al. Cytotoxic activity of Kingella kingae RtxA toxin depends on post-translational acylation of lysine residues and cholesterol binding. Emerg Microbes Infect 2018, 7, 178, doi:10.1038/s41426-018-0179-x.

11. Mazzone, A.; Ricevuti, G. Leukocyte CD11/CD18 integrins: biological and clinical relevance. Haematologica 1995, 80, 161–175.

12. Fine, D.H.; Patil, A.G.; Loos, B.G. Classification and diagnosis of aggressive periodontitis. J Periodontol 2018, 89 Suppl 1, S103–S119, doi:10.1002/JPER.16-0712.

13. Loe, H.; Brown, L.J. Early onset periodontitis in the United States of America. J Periodontol 1991, 62, 608–616.

14. Welch, R.A. Pore-forming cytolysins of gram-negative bacteria. Mol. Microbiol 1991, 5, 521–528.

15. Haubek, D.; Johansson, A. Pathogenicity of the highly leukotoxic JP2 clone of Aggregatibacter actinomycetemcomitans and its geographic dissemination and role in aggressive periodontitis. J Oral Microbiol 2014, 6, doi:10.3402/jom.v6.23980.

16. Brogan, J.M.; Lally, E.T.; Poulsen, K.; Kilian, M.; Demuth, D.R. Regulation of Actinobacillus actinomycetemcomitans leukotoxin expression: analysis of the promoter regions of leukotoxic and minimally leukotoxic strains. Infect. Immun. 1994, 62, 501–508.

17. Lally, E.T.; Golub, E.E.; Kieba, I.R.; Taichman, N.S.; Rosenbloom, J.; Rosenbloom, J.C.; Gibson, C.W.; Demuth, D.R. Analysis of the Actinobacillus actinomycetemcomitans leukotoxin gene. Delineation of unique features and comparison to homologous toxins. J. Biol. Chem. 1989, 264, 15451–15456.

18. Lally, E.T.; Kieba, I.R.; Sato, A.; Green, C.L.; Rosenbloom, J.; Korostoff, J.; Wang, J.F.; Shenker, B.J.; Ortlepp, S.; Robinson, M.K., et al. RTX toxins recognize a beta2 integrin on the surface of human target cells. J. Biol. Chem. 1997, 272, 30463–30469.

19. Brown, A.C.; Balashova, N.V.; Epand, R.M.; Epand, R.F.; Bragin, A.; Kachlany, S.C.; Walters, M.J.; Du, Y.; Boesze-Battaglia, K.; Lally, E.T. Aggregatibacter actinomycetemcomitans leukotoxin utilizes a cholesterol recognition/amino acid consensus site for membrane association. J. Biol. Chem. 2013, 288, 23607–23621, doi:10.1074/jbc.M113.486654.

20. Brown, A.C.; Koufos, E.; Balashova, N.; Boesze-Battaglia, K.; Lally, E.T. Inhibition of LtxA Toxicity by Blocking Cholesterol Binding With Peptides. Mol Oral Microbiol 2015, 10.1111/omi.12133, doi:10.1111/omi.12133.

21. Kinashi, T. Intracellular signalling controlling integrin activation in lymphocytes. Nat. Rev. Immunol. 2005, 5, 546–559, doi:10.1038/nri1646.

22. Li, N.; Yang, H.; Wang, M.; Lu, S.; Zhang, Y.; Long, M. Ligand-specific binding forces of LFA-1 and Mac-1 in neutrophil adhesion and crawling. Mol Biol Cell 2018, 29, 408–418, doi:10.1091/mbc.E16-12-0827.

23. Tohyama, Y.; Katagiri, K.; Pardi, R.; Lu, C.; Springer, T.A.; Kinashi, T. The critical cytoplasmic regions of the L/2 integrin in Rap1-induced adhesion and migration. Mol. Biol. Cell 2003, 14, 2570–2582, doi:10.1091/mbc.E02-09-0615 [doi];E02-09-0615 [pii].

24. Fabbri, M.; Fumagalli, L.; Bossi, G.; Bianchi, E.; Bender, J.R.; Pardi, R. A tyrosine-based sorting signal in the 2 integrin cytoplasmic domain mediates its recycling to the plasma membrane and is required for ligand-supported migration. EMBO J 1999, 18, 4915–4925, doi:10.1093/emboj/18.18.4915 [doi].

25. Caswell, P.T.; Norman, J.C. Integrin trafficking and the control of cell migration. Traffic 2006, 7, 14–21, doi:10.1111/j.1600-0854.2005.00362.x.

26. Fong, K.P.; Pacheco, C.M.; Otis, L.L.; Baranwal, S.; Kieba, I.R.; Harrison, G.; Hersh, E.V.; Boesze-Battaglia, K.; Lally, E.T. Actinobacillus actinomycetemcomitans leukotoxin requires lipid microdomains for target cell cytotoxicity. Cell. Microbiol. 2006, 8, 1753–1767, doi:10.1111/j.1462-5822.2006.00746.x.

27. Mahanonda, R.; Champaiboon, C.; Subbalekha, K.; Sa-Ard-Iam, N.; Yongyuth, A.; Isaraphithakkul, B.; Rerkyen, P.; Charatkulangkun, O.; Pichyangkul, S. Memory T cell subsets in healthy gingiva and periodontitis tissues. J Periodontol 2018, 89, 1121–1130, doi:10.1002/JPER.17-0674.

28. Li, Y.; Messina, C.; Bendaoud, M.; Fine, D.H.; Schreiner, H.; Tsiagbe, V.K. Adaptive immune response in osteoclastic bone resorption induced by orally administered Aggregatibacter actinomycetemcomitans in a rat model of periodontal disease. Mol Oral Microbiol 2010, 25, 275–292, doi:10.1111/j.2041-1014.2010.00576.x.

29. Lally, E.T.; Kieba, I.R.; Sato, A.; Green, C.L.; Rosenbloom, J.; Korostoff, J.; Wang, J.F.; Shenker, B.J.; Ortlepp, S.; Robinson, M.K., et al. RTX toxins recognize a β2 integrin on the surface of human target cells. J. Biol. Chem 1997, 272, 30463–30469.

30. Nygren, P.; Balashova, N.; Brown, A.C.; Kieba, I.; Dhingra, A.; Boesze-Battaglia, K.; Lally, E.T. Aggregatibacter actinomycetemcomitans leukotoxin causes activation of lymphocyte function-associated antigen 1. Cell. Microbiol. 2019, 21, e12967, doi:10.1111/cmi.12967.

31. Brown, A.C.; Boesze-Battaglia, K.; Balashova, N.V.; Mas Gomez, N.; Speicher, K.; Tang, H.Y.; Duszyk, M.E.; Lally, E.T. Membrane localization of the Repeats-in-Toxin (RTX) Leukotoxin (LtxA) produced by Aggregatibacter actinomycetemcomitans. PLoS One 2018, 13, e0205871, doi:10.1371/journal.pone.0205871.

32. Brown, A.C.; Boesze-Battaglia, K.; Du, Y.; Stefano, F.P.; Kieba, I.R.; Epand, R.F.; Kakalis, L.; Yeagle, P.L.; Epand, R.M.; Lally, E.T. Aggregatibacter actinomycetemcomitans leukotoxin cytotoxicity occurs through bilayer destabilization. Cell. Microbiol. 2012, 14, 869–881, doi:10.1111/j.1462-5822.2012.01762.x.

33. Lear, J.D.; Furblur, U.G.; Lally, E.T.; Tanaka, J.C. Actinobacillus actinomycetemcomitans leukotoxin forms large conductance, voltage-gated ion channels when incorporated into planar lipid bilayers. Biochim. Biophys. Acta 1995, 1238, 34–41.

34. Kumaresan, A.; Kadirvel, G.; Bujarbaruah, K.M.; Bardoloi, R.K.; Das, A.; Kumar, S.; Naskar, S. Preservation of boar semen at 18 degrees C induces lipid peroxidation and apoptosis like changes in spermatozoa. Anim. Reprod. Sci. 2009, 110, 162–171, doi:10.1016/j.anireprosci.2008.01.006.

35. Kirchhausen, T.; Macia, E.; Pelish, H.E. Use of dynasore, the small molecule inhibitor of dynamin, in the regulation of endocytosis. Methods Enzymol. 2008, 438, 77–93, doi:10.1016/S0076-6879(07)38006-3.

36. Hill, T.A.; Gordon, C.P.; McGeachie, A.B.; Venn-Brown, B.; Odell, L.R.; Chau, N.; Quan, A.; Mariana, A.; Sakoff, J.A.; Chircop, M., et al. Inhibition of dynamin mediated endocytosis by the dynoles--synthesis and functional activity of a family of indoles. J. Med. Chem. 2009, 52, 3762–3773, doi:10.1021/jm900036m.

37. Samuelsson, M.; Potrzebowska, K.; Lehtonen, J.; Beech, J.P.; Skorova, E.; Uronen-Hansson, H.; Svensson, L. RhoB controls the Rab11-mediated recycling and surface reappearance of LFA-1 in migrating T lymphocytes. Sci Signal 2017, 10, doi:10.1126/scisignal.aai8629.

38. Naslavsky, N.; Weigert, R.; Donaldson, J.G. Convergence of non-clathrin- and clathrin-derived endosomes involves Arf6 inactivation and changes in phosphoinositides. Molecular biology of the cell 2003, 14, 417–431, doi:10.1091/mbc.02-04-0053.

39. Balashova, N.; Giannakakis, A.; Brown, A.C.; Koufos, E.; Benz, R.; Arakawa, T.; Tang, H.Y.; Lally, E.T. Generation of a recombinant Aggregatibacter actinomycetemcomitans RTX toxin in Escherichia coli. Gene 2018, 672, 106–114, doi:10.1016/j.gene.2018.06.003.

40. Bretscher, M.S. On the shape of migrating cells--a ‘front-to-back’ model. J Cell Sci 2008, 121, 2625–2628, doi:10.1242/jcs.031120.

41. Bashkirov, P.V.; Akimov, S.A.; Evseev, A.I.; Schmid, S.L.; Zimmerberg, J.; Frolov, V.A. GTPase cycle of dynamin is coupled to membrane squeeze and release, leading to spontaneous fission. Cell 2008, 135, 1276–1286, doi:10.1016/j.cell.2008.11.028.

42. Pearse, B.M.; Bretscher, M.S. Membrane recycling by coated vesicles. Annu. Rev. Biochem. 1981, 50, 85–101, doi:10.1146/annurev.bi.50.070181.000505.

43. Aulik, N.A.; Hellenbrand, K.M.; Kisiela, D.; Czuprynski, C.J. Mannheimia haemolytica leukotoxin binds cyclophilin D on bovine neutrophil mitochondria. Microb. Pathog. 2011, 50, 168–178, doi:10.1016/j.micpath.2011.01.001.

44. Atapattu, D.N.; Albrecht, R.M.; McClenahan, D.J.; Czuprynski, C.J. Dynamin-2-dependent targeting of mannheimia haemolytica leukotoxin to mitochondrial cyclophilin D in bovine lymphoblastoid cells. Infect. Immun. 2008, 76, 5357–5365, doi:10.1128/IAI.00221-08.

45. Fabbri, M.; Di Meglio, S.; Gagliani, M.C.; Consonni, E.; Molteni, R.; Bender, J.R.; Tacchetti, C.; Pardi, R. Dynamic partitioning into lipid rafts controls the endo-exocytic cycle of the alphaL/beta2 integrin, LFA-1, during leukocyte chemotaxis. Molecular biology of the cell 2005, 16, 5793–5803, doi:10.1091/mbc.e05-05-0413.

46. Anderson, R.G. The caveolae membrane system. Annu. Rev. Biochem. 1998, 67, 199–225, doi:10.1146/annurev.biochem.67.1.199.

47. Oh, P.; McIntosh, D.P.; Schnitzer, J.E. Dynamin at the neck of caveolae mediates their budding to form transport vesicles by GTP-driven fission from the plasma membrane of endothelium. J. Cell Biol. 1998, 141, 101–114.

48. Parton, R.G.; Joggerst, B.; Simons, K. Regulated internalization of caveolae. J. Cell Biol. 1994, 127, 1199–1215.

49. Atapattu, D.N.; Czuprynski, C.J. Mannheimia haemolytica leukotoxin binds to lipid rafts in bovine lymphoblastoid cells and is internalized in a dynamin-2- and clathrin-dependent manner. Infect. Immun. 2007, 75, 4719–4727, doi:10.1128/IAI.00534-07.

50. Chinnapen, D.J.; Chinnapen, H.; Saslowsky, D.; Lencer, W.I. Rafting with cholera toxin: endocytosis and trafficking from plasma membrane to ER. FEMS Microbiol. Lett. 2007, 266, 129–137, doi:10.1111/j.1574-6968.2006.00545.x.

51. Zhen, Y.; Stenmark, H. Cellular functions of Rab GTPases at a glance. Journal of cell science 2015, 128, 3171–3176, doi:10.1242/jcs.166074.

52. Huotari, J.; Helenius, A. Endosome maturation. EMBO J. 2011, 30, 3481–3500, doi:10.1038/emboj.2011.286.

53. Rubino, M.; Miaczynska, M.; Lippe, R.; Zerial, M. Selective membrane recruitment of EEA1 suggests a role in directional transport of clathrin-coated vesicles to early endosomes. J. Biol. Chem. 2000, 275, 3745–3748, doi:10.1074/jbc.275.6.3745.

54. Damiano, J.S.; Cress, A.E.; Hazlehurst, L.A.; Shtil, A.A.; Dalton, W.S. Cell adhesion mediated drug resistance (CAM-DR): role of integrins and resistance to apoptosis in human myeloma cell lines. Blood 1999, 93, 1658–1667.

55. de la Fuente, M.T.; Casanova, B.; Moyano, J.V.; Garcia-Gila, M.; Sanz, L.; Garcia-Marco, J.; Silva, A.; Garcia-Pardo, A. Engagement of alpha4beta1 integrin by fibronectin induces in vitro resistance of B chronic lymphocytic leukemia cells to fludarabine. J Leukoc Biol 2002, 71, 495–502.

56. Balashova, N.; Dhingra, A.; Boesze-Battaglia, K.; Lally, E.T. Aggregatibacter actinomycetemcomitans leukotoxin induces cytosol acidification in LFA-1 expressing immune cells. Mol Oral Microbiol 2016, 31, 106–114, doi:10.1111/omi.12136.

57. DiFranco, K.M.; Kaswala, R.H.; Patel, C.; Kasinathan, C.; Kachlany, S.C. Leukotoxin kills rodent WBC by targeting leukocyte function associated antigen 1. Comp Med 2013, 63, 331–337.

58. Balashova, N.; Dhingra, A.; Boesze-Battaglia, K.; Lally, E.T. Aggregatibacter actinomycetemcomitans leukotoxin induces cytosol acidification in LFA-1 expressing immune cells. Mol Oral Microbiol 2015, 10.1111/omi.12136, doi:10.1111/omi.12136.

59. DiFranco, K.M.; Gupta, A.; Galusha, L.E.; Perez, J.; Nguyen, T.V.; Fineza, C.D.; Kachlany, S.C. Leukotoxin (Leukothera(R)) targets active leukocyte function antigen-1 (LFA-1) protein and triggers a lysosomal mediated cell death pathway. J. Biol. Chem. 2012, 287, 17618–17627, doi:10.1074/jbc.M111.314674.

60. Balashova, N.; Giannakakis, A.; Brown, A.C.; Koufos, E.; Benz, R.; Arakawa, T.; Tang, H.Y.; Lally, E.T. Generation of a recombinant Aggregatibacter actinomycetemcomitans RTX toxin in Escherichia coli. Gene 2018, 10.1016/j.gene.2018.06.003, doi:10.1016/j.gene.2018.06.003.

61. Platt, F.M.; d’Azzo, A.; Davidson, B.L.; Neufeld, E.F.; Tifft, C.J. Lysosomal storage diseases. Nat Rev Dis Primers 2018, 4, 27, doi:10.1038/s41572-018-0025-4.

62. DiFranco, K.M.; Johnson-Farley, N.; Bertino, J.R.; Elson, D.; Vega, B.A.; Belinka, B.A., Jr.; Kachlany, S.C. LFA-1-targeting Leukotoxin (LtxA; Leukothera(R)) causes lymphoma tumor regression in a humanized mouse model and requires caspase-8 and Fas to kill malignant lymphocytes. Leuk Res 2015, 39, 649–656, doi:10.1016/j.leukres.2015.03.010.

63. Kagedal, K.; Johansson, U.; Ollinger, K. The lysosomal protease cathepsin D mediates apoptosis induced by oxidative stress. FASEB J. 2001, 15, 1592–1594.

64. Kagedal, K.; Zhao, M.; Svensson, I.; Brunk, U.T. Sphingosine-induced apoptosis is dependent on lysosomal proteases. Biochem. J. 2001, 359, 335–343.

65. Kirkegaard, T.; Jaattela, M. Lysosomal involvement in cell death and cancer. Biochim. Biophys. Acta 2009, 1793, 746–754, doi:10.1016/j.bbamcr.2008.09.008.

66. Barcena-Uribarri, I.; Benz, R.; Winterhalter, M.; Zakharian, E.; Balashova, N. Pore forming activity of the potent RTX-toxin produced by pediatric pathogen Kingella kingae: Characterization and comparison to other RTX-family members. Biochim. Biophys. Acta 2015, 1848, 1536–1544, doi:10.1016/j.bbamem.2015.03.036.

67. Lear, J.D.; Karakelian, D.; Furblur, U.; Lally, E.T.; Tanaka, J.C. Conformational studies of Actinobacillus actinomycetemcomitans leukotoxin: partial denaturation enhances toxicity. Biochim. Biophys. Acta 2000, 1476, 350–362.

68. Friebe, S.; van der Goot, F.G.; Burgi, J. The Ins and Outs of Anthrax Toxin. Toxins (Basel) 2016, 8, doi:10.3390/toxins8030069.

69. Lemichez, E.; Bomsel, M.; Devilliers, G.; vanderSpek, J.; Murphy, J.R.; Lukianov, E.V.; Olsnes, S.; Boquet, P. Membrane translocation of diphtheria toxin fragment A exploits early to late endosome trafficking machinery. Mol. Microbiol. 1997, 23, 445–457.

70. O’Brien, D.P.; Hernandez, B.; Durand, D.; Hourdel, V.; Sotomayor-Perez, A.C.; Vachette, P.; Ghomi, M.; Chamot-Rooke, J.; Ladant, D.; Brier, S., et al. Structural models of intrinsically disordered and calcium-bound folded states of a protein adapted for secretion. Sci Rep 2015, 5, 14223, doi:10.1038/srep14223.

71. DiRienzo, J.M.; Tsai, C.C.; Shenker, B.J.; Taichman, N.S.; Lally, E.T. Monoclonal antibodies to leukotoxin of Actinobacillus actinomycetemcomitans. Infect. Immun. 1985, 47, 31–36.

72. Schneider, U.; Schwenk, H.U.; Bornkamm, G. Characterization of EBV-genome negative “null” and “T” cell lines derived from children with acute lymphoblastic leukemia and leukemic transformed non-Hodgkin lymphoma. Int. J. Cancer 1977, 19, 621–626.

73. Tsai, C.C.; Shenker, B.J.; DiRienzo, J.M.; Malamud, D.; Taichman, N.S. Extraction and isolation of a leukotoxin from Actinobacillus actinomycetemcomitans with polymyxin B. Infect. Immun. 1984, 43, 700–705.

74. Fine, D.H.; Furgang, D.; Kaplan, J.; Charlesworth, J.; Figurski, D.H. Tenacious adhesion of Actinobacillus actinomycetemcomitans strain CU1000 to salivary-coated hydroxyapatite. Arch Oral Biol 1999, 44, 1063–1076.

75. Diaz, R.; Ghofaily, L.A.; Patel, J.; Balashova, N.V.; Freitas, A.C.; Labib, I.; Kachlany, S.C. Characterization of leukotoxin from a clinical strain of Actinobacillus actinomycetemcomitans. Microb. Pathog. 2006, 40, 48–55, doi:10.1016/j.micpath.2005.10.005.

76. Nygren, P.; Balashova, N.; Brown, A.C.; Kieba, I.; Dhingra, A.; Boesze-Battaglia, K.; Lally, E.T. Aggregatibacter actinomycetemcomitans leukotoxin causes activation of lymphocyte function-associated antigen 1. Cell Microbiol 2018, 10.1111/cmi.12967, e12967, doi:10.1111/cmi.12967.

77. Altankov, G.; Grinnell, F. Fibronectin receptor internalization and AP-2 complex reorganization in potassium-depleted fibroblasts. Exp. Cell Res. 1995, 216, 299–309, doi:10.1006/excr.1995.1038.

78. Maldonado, R.; Wei, R.; Kachlany, S.C.; Kazi, M.; Balashova, N.V. Cytotoxic effects of Kingella kingae outer membrane vesicles on human cells. Microb. Pathog 2011, 51, 22–30, doi:S0882-4010(11)00057-X [pii];10.1016/j.micpath.2011.03.005 [doi].

79. Parker, H.; Chitcholtan, K.; Hampton, M.B.; Keenan, J.I. Uptake of Helicobacter pylori outer membrane vesicles by gastric epithelial cells. Infect. Immun 2010, 78, 5054–5061, doi: IAI.00299-10 [pii];10.1128/IAI.00299-10 [doi].

80. Benz, R.; Janko, K.; Boos, W.; Lauger, P. Formation of large, ion-permeable membrane channels by the matrix protein (porin) of Escherichia coli. Biochim. Biophys. Acta 1978, 511, 305–319.

81. Dutta, D.; Donaldson, J.G. Search for inhibitors of endocytosis: Intended specificity and unintended consequences. Cell Logist 2012, 2, 203–208, doi:10.4161/cl.23967.

82. Kilsdonk, E.P.; Yancey, P.G.; Stoudt, G.W.; Bangerter, F.W.; Johnson, W.J.; Phillips, M.C.; Rothblat, G.H. Cellular cholesterol efflux mediated by cyclodextrins. J. Biol. Chem. 1995, 270, 17250–17256.

83. Larkin, J.M.; Brown, M.S.; Goldstein, J.L.; Anderson, R.G. Depletion of intracellular potassium arrests coated pit formation and receptor-mediated endocytosis in fibroblasts. Cell 1983, 33, 273–285.

